# Intracellular injection of brain extracts from Alzheimer’s disease patients trigger unregulated Ca^2+^ release from intracellular stores that hinders cellular bioenergetics

**DOI:** 10.1101/2022.04.29.490088

**Authors:** Anna Pensalfini, Abdul Rahim Umar, Charles Glabe, Ian Parker, Ghanim Ullah, Angelo Demuro

**Affiliations:** Department of Molecular Biology and Biochemistry, University of California, Irvine, CA; Department of Physics, University of South Florida, Tampa, FL; Department of Neurobiology and Behavior, University of California, Irvine, CA; Department of Physiology and Biophysics, University of California, Irvine, CA

## Abstract

Strong evidence indicates that amyloid beta (Aβ) inflicts its toxicity in Alzheimer’s disease (AD) by promoting uncontrolled elevation of cytosolic Ca^2+^ in neurons. We have previously shown that synthetic Aβ42 oligomers stimulate abnormal intracellular Ca^2+^ release from the endoplasmic reticulum stores, suggesting that a similar mechanism of Ca^2+^ toxicity may be common to the endogenous Aβs oligomers. To investigate this possibility, we use human postmortem brain extracts from control and AD-affected patients and test their ability to trigger Ca^2+^ fluxes when injected intracellularly into *Xenopus* oocytes. Immunological characterization of samples from AD patients revealed elevated content of soluble Aβ oligomers, detected by the conformation-dependent OC-antibody, whereas no immunoreactivity was detected in the normal samples. Intracellular injection of brain extracts from control patients failed to trigger detectable changes in intracellular Ca^2+^. Conversely, brain extracts from AD patients triggered Ca^2+^ events consisting of local and global Ca^2+^ fluorescent transients rising within few seconds after injection and persisting for several seconds. Pre-incubation of brain extracts with the conformation specific OC antibody completely suppressed brain extract ability to trigger cytosolic Ca^2+^ events. Comparison of the elementary events triggered by brain extracts and synthetic Aβ42 oligomer showed comparable temporal evolution and amplitudes to events triggered by direct injection of IP3. Moreover, bath application of caffeine reversibly inhibited local and global Ca^2+^ signals in all the samples confirming the involvement of Ca^2+^ release from the ER. Analysis of the recorded Ca^2+^ fluorescence signals by computational modeling allowed quantification of the IP3 and Ca^2+^ generated by each sample. The model further shows that the abnormal increase of Ca^2+^ and IP3 may affect mitochondrial bioenergetics. These results, supports the hypothesis that endogenous amyloid oligomer contained in neurons of AD-affected brains may represent the toxic agents responsible for neurons malfunctioning and death, associated with the disruption of neuronal Ca^2+^ homeostasis.

## Introduction

Progressive disruption of neuronal Ca^2+^ homeostasis is one of the leading mechanisms of action of the amyloidogenic proteins Aβ as the toxic species in the etiology of Alzheimer’s disease (AD) [1-3]. Increase in intracellular content of oligomeric Aβs is believed to play a major role in the early phase of AD as their intracellular rise strongly correlates with the symptoms and have been proved to be predictive of cognitive status in AD affected patients [4-6].

Previous results from our lab support the notion that intracellular as well as extracellular interaction of physiological Aβ42 oligomers with cell membranes, promote cytosolic Ca^2+^ rise to a toxic level [7-9]. Pharmacological and computational studies reveal that application of synthetic Aβ42 oligomers to different cells type, induce a G-protein-mediated stimulation of IP3 production, triggering intracellular Ca^2+^ fluxes leading to a disruption of intracellular Ca^2+^ homeostasis. [7, 8, 10, 11]

These findings led us to propose that abnormal stimulation of IP_3_ overproduction triggered by Aβ42 oligomers may contribute importantly to Ca^2+^ signaling disruptions and neurotoxicity in AD-affected neurons. While there is a widespread agreement on the toxic properties Aβ42 oligomers, most of this knowledge comes from the use of synthetic Aβs peptides, aggregated in vitro using distinct protocols generating ambiguity weather endogenous Aβs oligomers may carry out similar effects on neurons [2, 12-14].

The recent ability to obtain human brain extract from AD-affected brain, containing high levels of endogenous Aβ aggregates, offer the opportunity to validate these previous findings by investigating their ability to disrupt intracellular Ca^2+^ signaling, a condition observed in neurons of AD-affected brains. Here we investigate the Ca^2+^ dependent toxicity of selected brain extracts, chosen from postmortem brain samples that have been previously characterized by immunohistochemistry and found to contain high level of OC-positive oligomers. Fluorescence Ca^2+^ imaging experiments were performed using seven different samples of brain extracts: three from normal brain (N1-B11, N2-B11 and N3-B11) and four from AD affected samples (AD1-B11,

AD2-B11 and AD3-TEC and AD3-B11) [4, 5]. Experiments were performed by intracellular microinjection of 10 nl of these samples into *Xenopus* oocytes — loaded with Ca^2+^ sensitive dye — and their ability to trigger cytosolic Ca^2+^ signal was tested. Injections of brain extracts from normal patients evoked little or non-detectable fluorescent Ca^2+^ signals. On the contrary, intracellular injection of extracts from AD brains consistently triggered cytosolic Ca^2+^ elevations. These Ca^2+^ events consist of both local and global Ca^2+^ signals, and showed strong similarity with those evoked after injection of synthetic Aβ42 oligomers and intracellular IP3 release [15, 16]. Blockage by caffeine and OC-antibody support the ability of endogenous Aβ aggregates contained in the AD brains extracts to trigger cytosolic Ca^2+^ release by uncontrolled activation of IP3Rs [17-19]. Next, we used computational modeling to quantify the corresponding cytosolic increase in IP3 and Ca^2+^ and the resulting disruption of the normal cell bioenergetics. Overall, our results support the notion that early intracellular rise in Aβ aggregates seen in AD-affected neurons may inflict neuronal cytotoxicity by disrupting intracellular Ca^2+^ homeostasis and cellular bioenergetics.

## Methods

### Oocyte preparation and microinjection

We used *Xenopus* oocytes as a model cell preparation, as their large size enables intracellular microinjection of samples [20]. Experiments were performed on stage VI oocytes, injected before imaging with fluo-4-dextran (MW 10,000 D; kD for Ca^2+^ about 3 mM) to a final intracellular concentration of ∼40 μM. Oocytes were then placed animal hemisphere down in a chamber whose base is formed by a fresh, ethanol-washed microscope cover glass (Fisherbrand, type-545-M), and bathed in a Ca^2+^ free Ringer’s solution (composition in mM: NaCl, 110; KCl, 2; HEPES, 5; at pH 7.2.). A gravity-fed superfusion system allows exchange of the Ringer’s solution. Oocytes were imaged at room temperature by wide-field fluorescence microscopy using an Olympus inverted microscope (IX 71) (Figure 1A) equipped with a 60X oil-immersion objective, a 488 nm solid state laser for fluorescence excitation and a ccd camera (Cascade 128+: Roper Scientific) for imaging fluorescence emission at frame rates of 10 to 30 s-1. Changes in fluorescence intensity was imaged within a 40 × 40 μm region in the animal hemisphere of the oocyte and measurements are expressed as a ratio (ΔF/Fo) of the change in fluorescence at a given region of interest (ΔF) relative to the resting fluorescence at that region before stimulation (Fo). Mean values of Fo were obtained by averaging over several frames before stimulation using MetaMorph (Molecular Devices). Fluorescent traces were exported to Microcal Origin version 6.0 (OriginLab,Northamptom, MA, USA) for analysis and graphing. Microinjection of 10 nl of brain extract samples into oocytes was performed using a Drummond nanoinjector mounted on a hydraulic micromanipulator. A glass pipette (tip diameter of 8-10 μm) was filled with samples and inserted vertically down across the entire oocyte to a pre-established position with the tip positioned 6 to 8 μm inward from the plasma membrane and centered within the image field.

**Figure 1).**
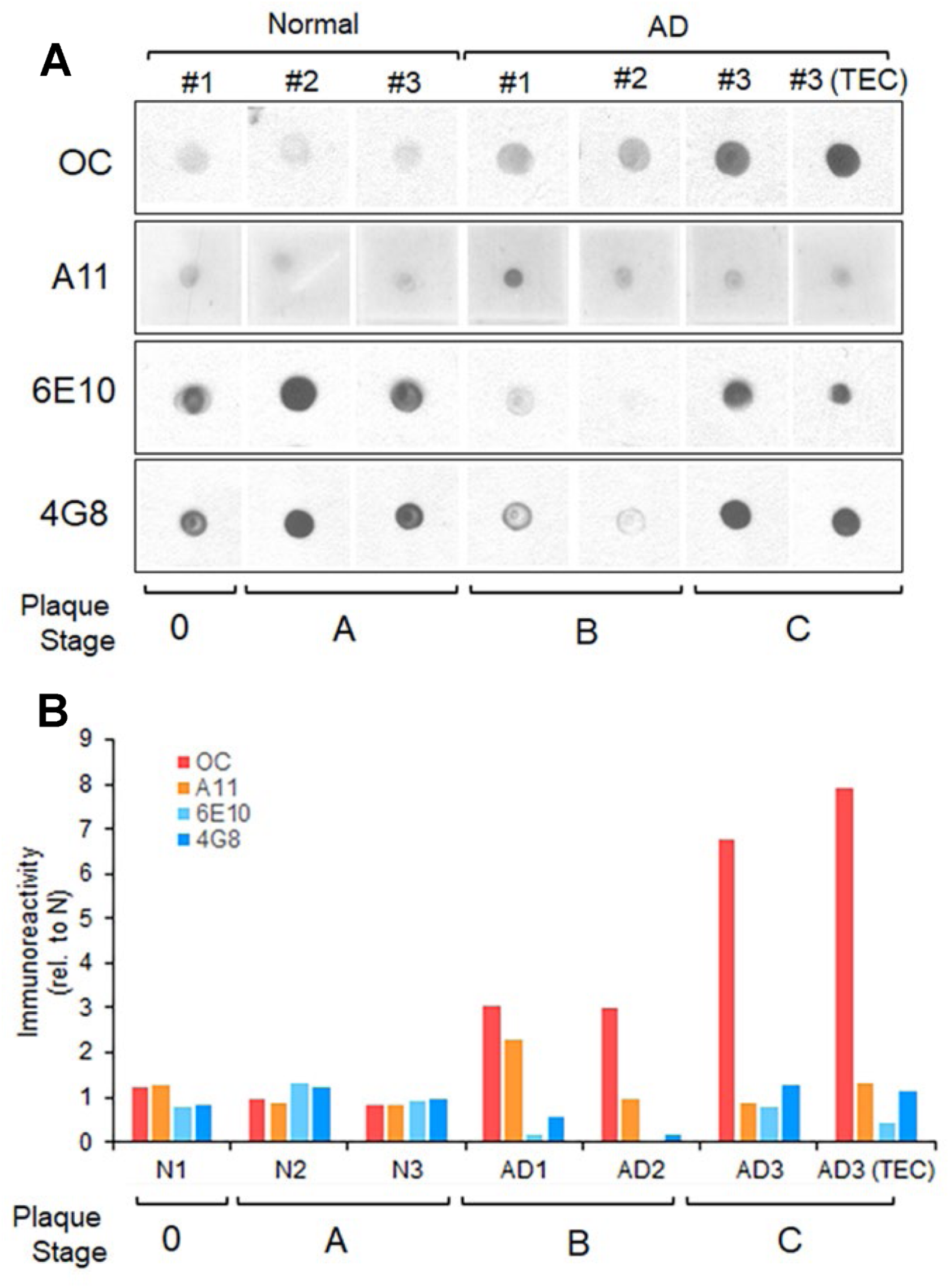
Immunological characterization of human AD brains extract reveals high content of OC positive fibrillar oligomers. Dot blot analysis, and respective quantification, of human soluble PBS fractions from the B11 and TEC regions of normal and AD patients probed with OC, A11, 6E10 and 4G8 antibodies. OC positive fibrillar oligomers, and not A11 positive prefibrillar oligomers, are increased in AD brain sample with increasing plaque stage pathology. The apparent decrease in 6E10 and 4G8 immunoreactivities in the AD patients with intermediate plaque pathology (stage B) suggests that at this stage Aβ undergoes a conformational change that is not recognized by 6E10 or 4G8, but that can be readily detected by OC.

### Patient brain tissue selection

Frozen brain tissue was obtained from the UCI Alzheimer Disease Research Center (ADRC). Subjects enrolled in the ADRC were given the MMSE. As a standard protocol for ADRC autopsy cases, Braak & Braak neurofibrillary tangle and plaque staging was evaluated [21]. Our objective was to test brains samples with high content of OC positivity to antibody. We selected tissues from the frontal cortex (Brodmann’s Area11; B11) from two different AD-affected individuals, and from a third individual from which both B11 and trans-entorhinal cortex (TEC) samples where available (Table 1). These subjects were widely characterized in our previous studies for their immunoreactivity with various conformation-dependent antibodies, such as OC and M78, which correlated with cognitive decline, tangle stage and plaque pathology, as well as with early intracellular/intranuclear aggregates build up at intermediate stages of plaque pathology (plaque stage A-B), respectively [4, 5]. Table 1 lists the clinical and pathological details of the cases used in this study. The stock solution for all the brain extract was estimated to be 1μg/ml for Aβ contents in PBS. For comparison with our previous investigations using synthetic Aβ42 oligomers, most of the experiment reported here are performed injecting 1μg/ml. except for the experiments where OC antibody were used to inhibit endogenous Aβs activity where they were used with OC of 0.5 μg/ml.

**Table 1:**
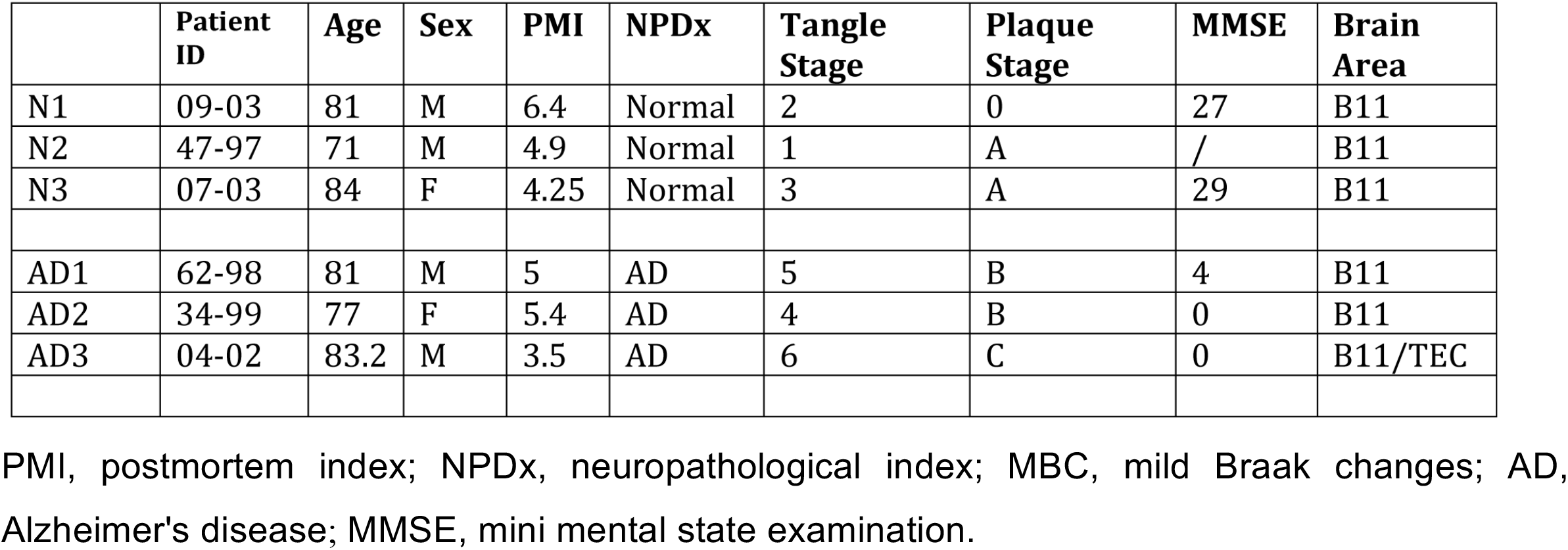
Brain extracts tested

### Human brain sample preparation and dot blot analysis

Frozen tissues from the B11 area or trans-entorhinal cortex (TEC) were processed as previously described [5]. Briefly, the brains were weighted, diced, and homogenized in ice-cold PBS (4:1 PBS volume/brain wet weight), 0.02% NaN3, pH 7.4, supplemented with protease inhibitor cocktail. The samples were ultracentrifuged at 100,000 × G for 1 hr at 4°C. The PBS soluble fraction was collected, aliquoted and stored at – 80°C for future testing.

For dot blot analysis, 2 μg of samples was spotted in duplicate onto nitrocellulose membrane and allowed to air dry. After blocking the non-reactive sites with 10% nonfat dry milk in low-Tween TBS-T, (20 mM Tris, 137 mM NaCl, 0.01% Tween 20 pH 7.6), the blots were incubated overnight at 4°C in primary antibody at the following dilutions: OC 1:10,000, A11 1:2000, 6E10 (Covance, Princeton, NJ) 1:2000. The blots were then incubated with goat-α-rabbit (for OC and A11) and goat-α-mouse HRP conjugated secondary antibody (Jackson Immune Research 1:12000), washed three times for 5 min, followed by ECL detection. Stock solution for OC was 0.4 μg/ml in PBS solution.

### Computational Methods

Equations modeling cytosolic Ca^2+^ dynamics and mitochondrial function, and numerical methods are described in detail in Supplementary Information Text.

## Results

### Brain extract from AD patients display high content of OC-positive Aβs

Of relevant interest in understanding the role of Aβs oligomers in the etiology of AD, is to uncover the prevailing type of aggregates that more closely associate with the development of the symptoms, and the associated molecular mechanisms [22]. Previous work from various labs including ours, has demonstrated that the time dependent accumulation of intracellular Aβs oligomers is strictly correlate with the progression of the symptoms over time. In addition, intracellular Ca^2+^ rise observed in neurons during aging has been linked to the action of a specific oligomeric forms of Aβs.

In this work we selected samples of postmortem brain extracts form three normal and three AD-affected patients which were screened for their affinity to selected antibodies for different types of Aβ aggregates [4, 5]. Figure 1A shows dot blot analysis of human soluble PBS fractions probed with OC, A11, 6E10 and 4G8 antibodies. OC positive oligomers, and not A11, 6E10 nor 4G8 immunoreactivity is increased in AD brain samples compare to normal samples. In Figure 1B, a comprehensive plot shows the calculated fold changes in immunoreactivity of each antibody. The plot shows that very low content of OC positive Aβs oligomers was found in samples N1-B11, N2-B11 and N3-B11 from normal brains, whereas high content of OC positive oligomers was detected in all the samples from AD affected brains AD1-B11, AD2B11 with particular high immunoreactivity in AD3-B11/TEC samples.

### Intracellular injection of brain extracts induces local and global cytosolic Ca^2+^ fluxes

To examine the ability of human brain extract to trigger Ca^2+^ mobilization from intracellular stores, we use fluorescent Ca^2+^ imaging with the *Xenopus* oocyte as a model cell system. The experiments described here follow similar approaches we previously applied to investigate synthetic Aβ42 oligomers Ca^2+^ toxicity [8]. The stock solution of brain extracts used in this work have been estimated to contain up to 1 μg/ml of Aβ in PBS solution. Intracellular injection of 10 nl samples at this concentration triggered fast and ample fluorescent responses. However, we observed that injection of 10 nl of lower concentrations, down to 0.3 μg/ml dilutions retain the ability to consistently triggered Ca^2+^ signaling (data not shown).

Intracellular microinjection of 10 nl samples from AD brain extracts, evoked potent cytosolic Ca^2+^ mobilization of different spatiotemporal patterns ranging from repetitive local short events such as blips and puffs to global repetitive Ca^2+^ waves covering the entire image field. Conversely, injection of 10 nl of brain extracts from normal individuals failed to promote either local or global Ca^2+^ fluxes. In Figure 2, we show results from selected series of single experiments depicting the spatiotemporal evolution of the Ca^2+^-dependent fluorescent signals evoked by injection of different brain extracts. In Figure 2A, the top trace depicts the Ca^2+^-fluorescent signal evoked in response to intracellular injection of 10 nl of AD3-TEC sample at the time indicated by the arrow in the line-scan. Below is the corresponding line-scan (kymograph) image, showing temporal evolution of the fluorescent signal originated by measuring changes in fluorescence intensity along a single line scan (25 μm long and 5 μm wide) properly positioned above the centroid of the fluorescent signal. The panel, display its evolution over time as a pseudo-colored representation where warmer color indicates increase in fluorescence signals intensities. Fluorescent transient with different amplitudes and duration are spotted along the line-scan reminiscent of the hierarchical organization of the of the IP3 mediated Ca^2+^ signaling. Figure 2B and 2C illustrate the corresponding fluorescent in response to injection of AD3-B11 and AD1-B11 samples showing large variability in time and amplitudes of the fluorescent responses. This variability was shared among all the tested AD-affected samples, independently from the donor or brain region (B11 or TEC). Figure 2D shows a typical response obtained after intracellular injection of 10 nl of samples from normal brain, which consistently failed to give detectable fluorescent responses.

**Figure 2).**
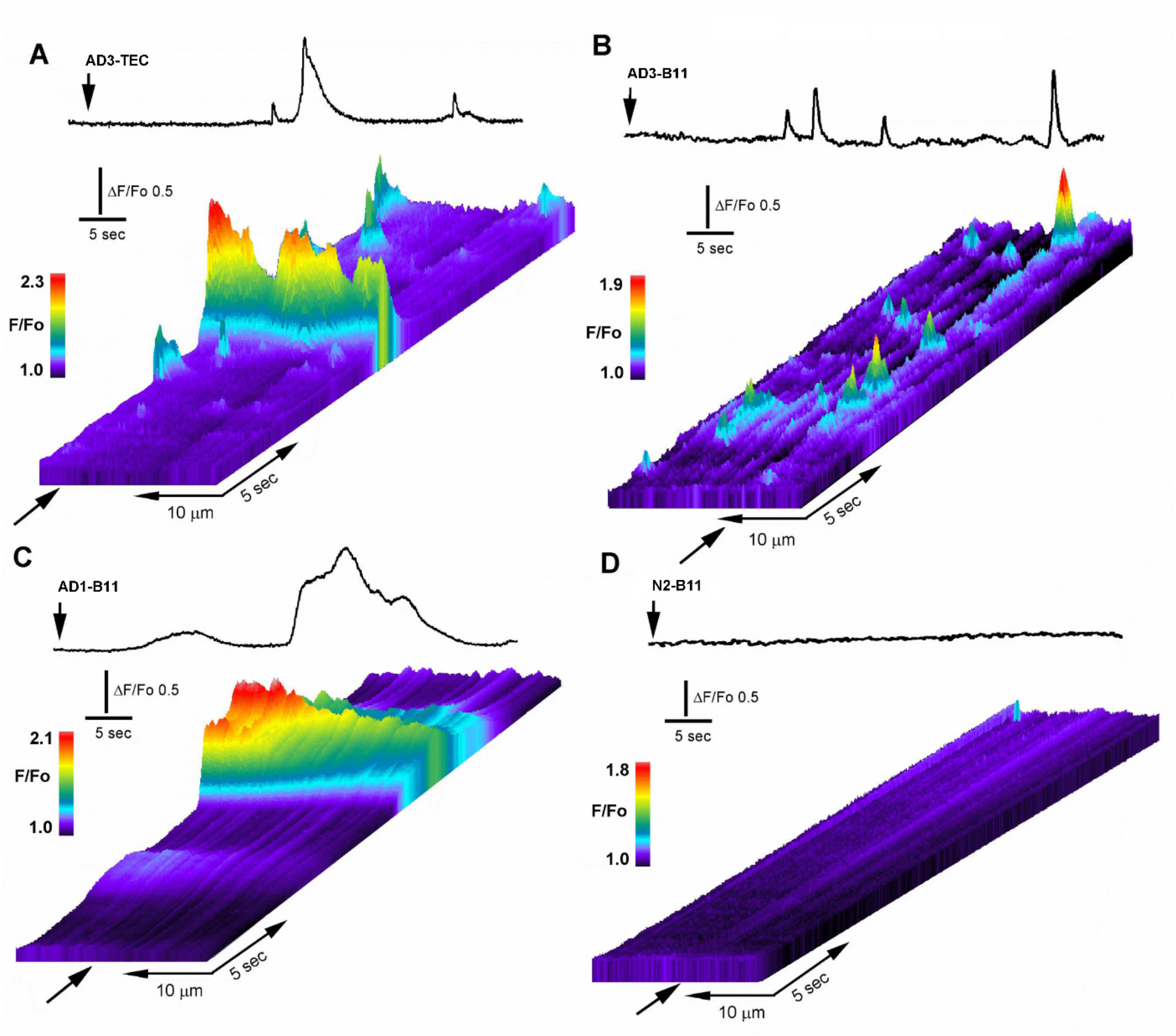
Intracellular injection of OC-positive brain extracts trigger local and global intracellular Ca^2+^ transients. **A**, Line-scan (kymograph) images illustrating spatiotemporal pattern of fluorescence Ca^2+^ signals recorded from an oocyte injected with 10 nl of AD3-TEC. Top trace shows fluorescence signal monitored from a small region along the line-scan positioned as marked by the arrow in the line-scan image. Brain extract was injected at the time indicated by the vertical arrow. Increasing fluo-4 pseudo-ratio signals are represented by warmer colors as depicted by the color bar and by increasing height at each pixel. Top traces in **B** and **C**, are respective fluorescence traces obtained by imaging oocytes injected with AD3-B11 and AD2-B11 respectively. Fluorescent traces are generated by averaging fluorescent intensity in a region of interest positioned on top footprint of the image over time. In **D**, trace show average fluorescent signal measured across the image field and kymograph image of the fluorescent background in an oocyte injected with brain extract from the normal patient (N1-B11).

In Figure 3A a selected number of fluorescent traces (black) from 23 different injections including a proportional number of traces from each sample of AD-affected brain extracts are shown. Traces are aligned to the first initial response for each injection and are not indicative of time delay of the fluorescence response after injection. Control experiment using sample from normal individual, regularly failed to evoke fluorescent response. Selected fluorescent traces obtained from experiment using different samples from control brains are superimpose and plotted together in Figure 3B. The overall results obtained in this series of experiments in outlined in figure 3D, where the number of responding cells from all the experiments are reported as percent of the total cells tested. In all the experiments performed, independently from the gender of the donor and the specific area of the brain from which the sample originate, samples from AD patients consistently triggered fluorescent responses whereas comparable samples from normal donor of equivalent age consistently failed to evoke significant fluorescent responses. Only a single response consisting of a single localized brief fluorescent transient (puff) was detected throughout the imaged field that we attributed to endogenous spontaneous Ca^2+^ signaling. We applied similar approach for screening all the trials performed and fluorescent responses consisting in a minimum of two separated sites with repetitive transients at each site were classified as active responses.

**Figure 3).**
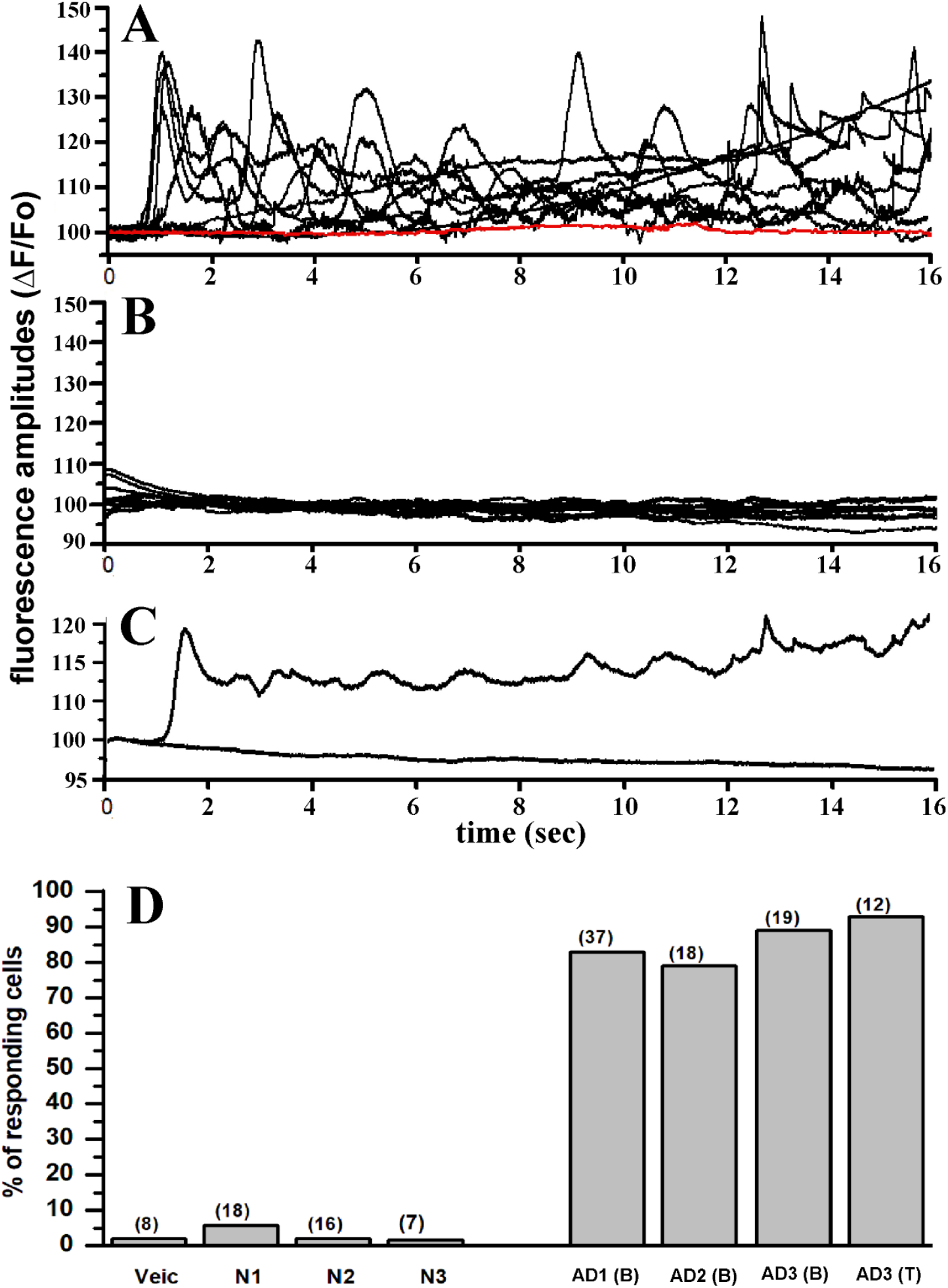
Intracellular injection of brain extracts from AD patients evoked large and consistent intracellular Ca^2+^ signals which are absent upon injection of brain extracts from non-AD patients. **A)** Superimposed traces represent fluorescent signals recorded from 10 different oocytes in response to intracellular injection with (10 nl) of OC-positive brain extract from patient AD1(B) (see Table 1 for description). Red trace indicates fluorescent background in non-injected oocyte. **B**) Superimposed fluorescent traces from 10 different oocytes recorded after injection with brain extracts from non-AD patient. **C**) Plot shows averaged fluorescent traces from **A** and **B. D)** Bar graph reporting the number of oocytes showing activation of fluorescence Ca^2+^ signals after intracellular injection of 10 nl of : PBS (veicle), N1, N2, N3 brain extract from normal patients (see Table 1), and brain extract from AD affected patients – AD1-B11, AD2-B11, AD3-B11 and AD3-TEC. Each column represents the number of responding oocytes expressed as percentage of the total number of oocytes tested for each group. Number of oocytes for each group in indicated by the number on top of each column.

### Conformation dependent OC-antibody strongly inhibit Ca^2+^ mobilization by AD-extracts

To further strength our hypothesis that the soluble Aβs oligomers contained in the AD-affected samples were responsible for the Ca^2+^ dependent fluorescent transients observed after injection, we made the use of the conformation specific OC antibody, with the intent of neutralizing the ability of endogenous Aβ. To this aim, we performed a series of experiments where, samples from AD brains were pre-incubated for 24 hours with OC-antibody prior to injection. In parallel, equivalent aliquots of corresponding samples were incubated without addition of antibody. In Figure 4A top-left, the trace depicts the temporal evolution of the fluorescent signal in response to intracellular injection of 10 nl AD3-TEC sample at the time indicated by the arrow. In this specific experiment, within 10 seconds after injection a robust, fast rising fluorescent response was observed which covered a large portion of the imaged field (40×40 μm^2^). Trace was obtained by measuring the average intensity within a region of interest covering the footprint of the fluorescent signal. This time delay in the fluorescent response was very common in all the experiments and we never observed fluorescent response with a delay longer than 30-40 secs. We then performed parallel experiment using the corresponding sample which have been previously incubated with OC antibody. Top-right trace in Figure 4A depicts the averaged intensity of the fluorescent signal recorded after injection of 10nl of AD3-TEC+OC. The sample was injected at the time indicated by the arrow and with ∼ 30 second delay a pulse of UV was applied as to ensure functional capability of the IP3 signaling system. Similar experiment for sample AD3-B11 and AD2-B11 are reported in the middle two traces and bottom two traces of Figure 4A respectively. The cumulative result of this set of experiment is reported in Figure 4B. Each column in the plot represents the number of responding cells as percentage of the total cell tested as indicated on top of each column.

**Figure 4).**
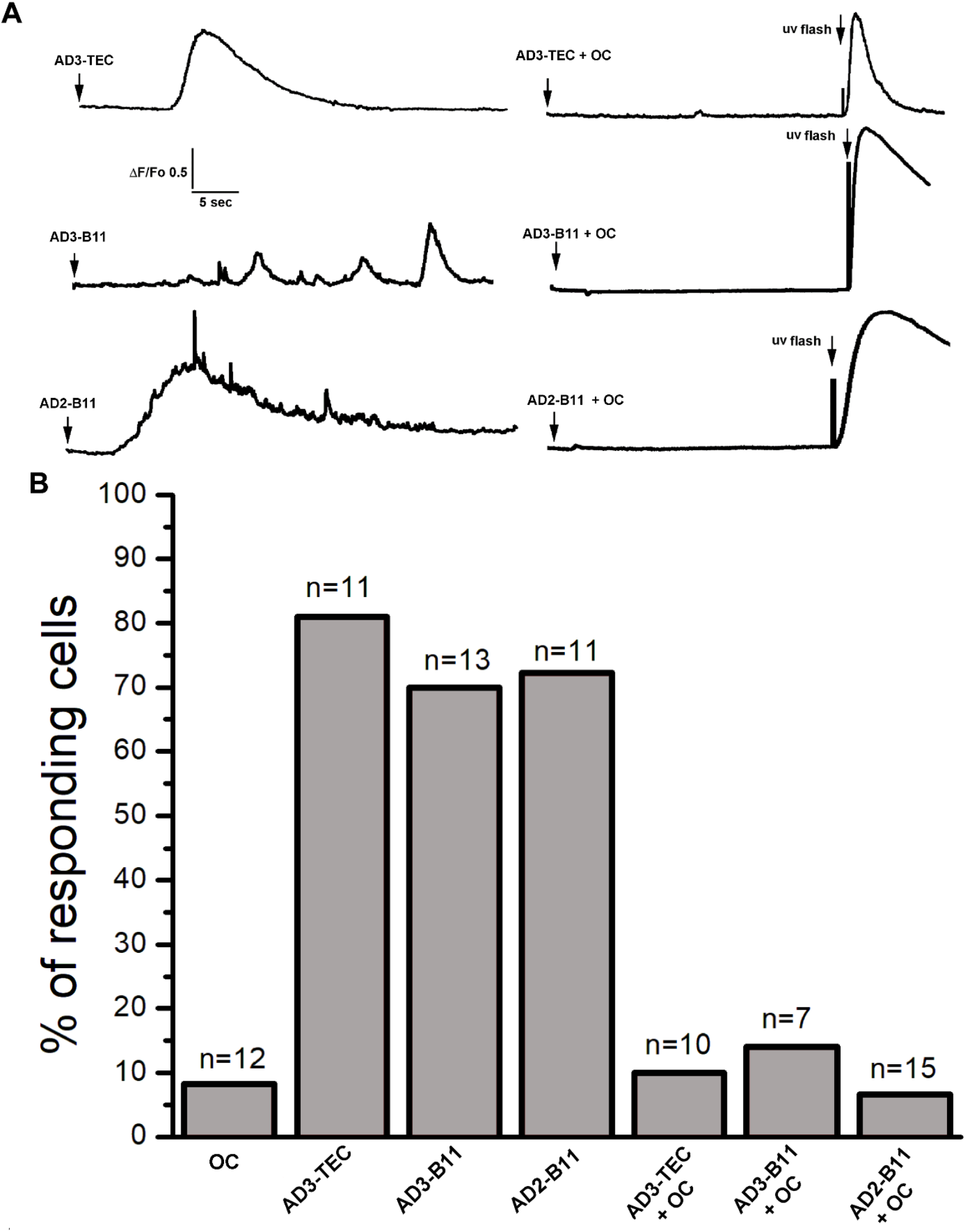
Incubation of AD brain extracts with OC antibody strongly inhibits their ability to activate Ca^2+^ signals. **A)** Top-left trace shows a Ca^2+^ fluorescent transient triggered by injection of 10 nl of AD1-TEC (Aβ 0.5 μg/ml) at the time indicated by the arrow. Ca^2+^ response was abolished by preincubation of AD1-TEC sample with OC antibody (OC 0,2 μg/ml ; Aβ 0.5 μg/ml). At the end of the recording a UV flash was given to trigger photorelease of IP_3_ as a control. The middle traces show fluorescence Ca^2+^ signals recorded from oocytes injected with AD1-B11 and corresponding fluorescence record obtained after injection of pre-incubated AD1-B11 with OC antibody. Bottom traces are obtained from oocyte injected with AD2-TE (left), and after incubation with OC antibody (right). B) Cumulative results showing percent of responding cells for each sample. Columns, represents numbers of oocytes responding to intracellular injection from samples indicated at the bottom of the graph. The data are expressed as percentage of total number of oocytes tested for each group. n at the top of each column represent the total number of oocytes tested. Oocytes were injected with OC antibody alone as control.

### Ca^2+^ fluxes triggered by endogenous Aβs is initiated by IP3-mediated intracellular liberation

The close resemblance between the temporal evolution of the fluorescence responses triggered by intracellular injection of the AD-samples used in this investigation and the IP3-mediated Ca^2+^ signaling suggests that intracellular AD brain extracts promote the liberation of Ca^2+^ from the ER stores via opening of IP_3_Rs. Moreover, following similar approach, we have previously reported that intracellular injection of synthetic Aβ42 oligomers into Xenopus oocytes triggers cytosolic Ca^2+^ release through a mechanism involving IP_3_ overproduction. In Figure 5, we show a comparative analysis of localize Ca^2+^ events triggered by endogenous Aβs (Figure 5A), synthetic Aβ42 oligomers (Figure 5B), and by injection of 10 nM IP_3_ (Figure 5C). In these analyses only selected experiments where local Ca^2+^ events were triggered are included. In Figure 5A, the top image is a kymograph illustration of puff-like events occurring at the same location, promoted by intracellular injection (10 nl) of the endogenous Aβ oligomers. Image is generated from the measurement of fluorescence along a line (*y*-axis; 5 μm long and 2 μm wide) positioned above the puff’s footprint in the video record, with time running left to right along the *x*-axis. Increasing fluorescent signals is represented by warmer colors as indicate by the color bar and by increasing height of each pixel. Below, the left plot is generated by the overlapping of 10 randomly selected traces aligned to the initial rise of the fluorescent trace. Inset is generated by averaging all the puffs traces. Center and right plots in Figure 5A show distributions of puff durations (measure at half-maximal amplitude) and maximum amplitudes obtained by measurement of 287 puffs from 6 oocytes from experiments carried out with different brain extracts.

**Figure 5).**
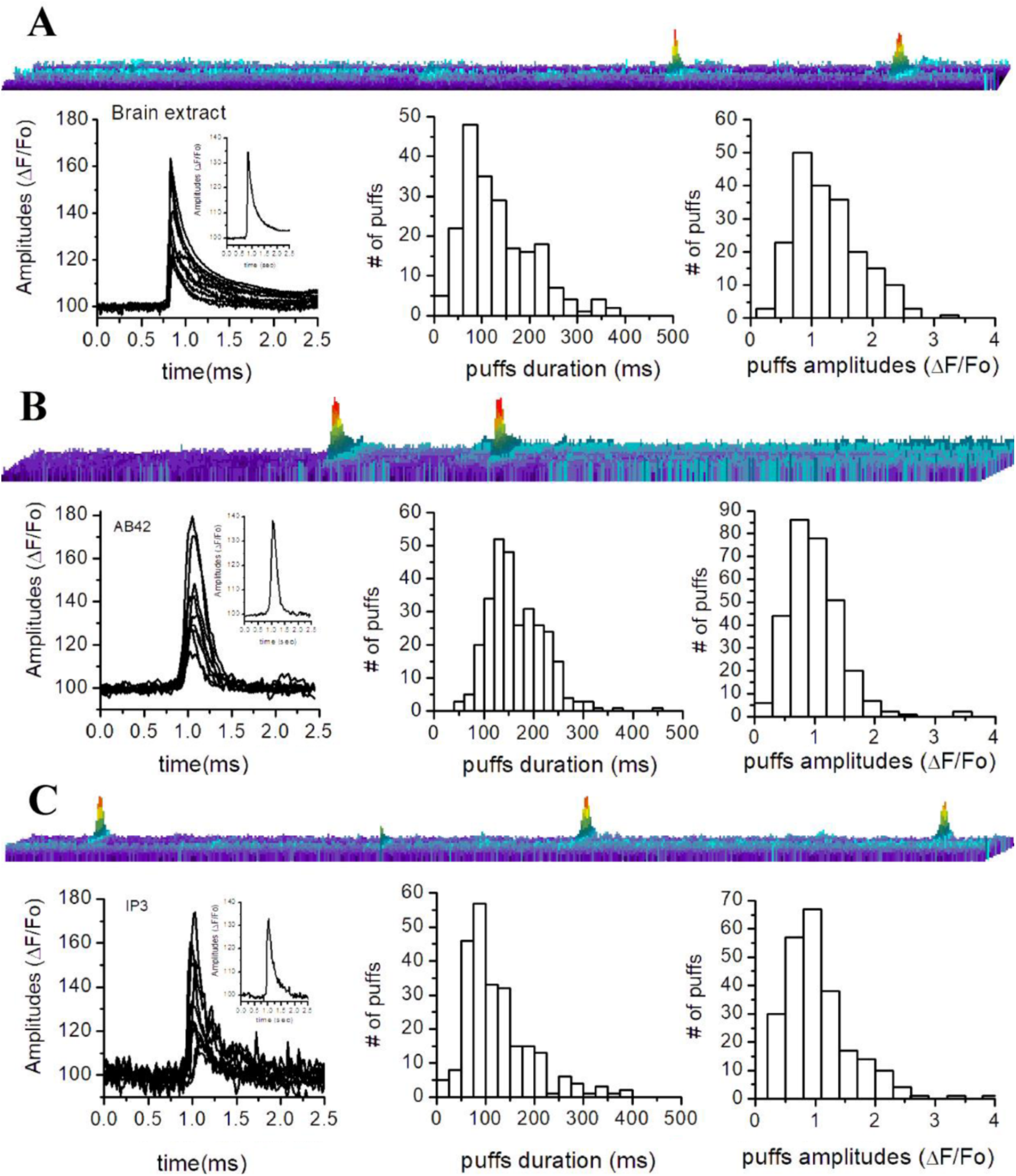
Amplitude and temporal evolution of local Ca^2+^ signals triggered by Intracellular injection of brain extracts from AD patients closely resemble those triggered by synthetic Aβ42 oligomers and by IP3. **A)** Top image is a line-scan (kymograph) representation of local intracellular Ca^2+^ events, evoked by intracellular injection (10 nl) of AD1-TEC. Below, left plot show superimposed traces from 10 different events. Inset is the resulting averaged profile. Center and right plots show distributions of events duration (mean duration 132.30+/-5.1 ms) and amplitudes (mean amplitude F/Fo 1.378+/-0.058) measure from 208 events in 8 oocytes. Data are from oocytes injected with either AD3-B11(n=2), AD3-TECA (n=3) or AD2-B11 (n=3). **B)** Results from parallel experiments injecting 10 nl of synthetic of Aβ42 oligomers (1μg/ml). Top image is a line-scan depicting fluorescent intensity over time. Left plot is generated by superimposing traces from 10 selected events and inset is their resulting averaged profile. Center end left plots are distributions of events durations (mean duration 165.06+/-3.3 ms) and amplitudes (mean amplitude F/Fo 100.3+/-2.58) measured from 297 events in 4 oocytes. **C)** Top image depicts a line-scan measurement of fluorescent signal over time after injection of 10 nl of 30 nM solution of IP_3_. The plot below show superimposed traces of 10 puffs and inset is the corresponding averaged profile. On the right are distributions of event durations (mean duration 124.06+/-4.9 ms) and amplitudes (mean amplitude F/Fo 1.11+/-0.045) measured from 243 puffs from 4 oocytes.

Figure 5B shows analysis of fluorescent events evoked by intracellular injection of 10 nl of synthetic Aβ42 oligomers (1 μg/ml). Top image illustrates repetitive puffs occurring at the same location in the image field. Below, the left plot is generated by the overlapping traces from 10 selected puffs. The trace in the inset is generated averaging the 10 traces in main plot. Center and right plots are the distribution of the measured puff durations and amplitudes. Figure 5C illustrates comparative analyses of the fluorescent transients generated directly by intracellular injection of 10 nl of IP3 (10 nM). Top image displays three fluorescent transients occurring at the same location, closely resembling the local transients observed when synthetic and endogenous

Aβ oligomers are injected. Overlapping of the puff-like profiles is shown in left plot and inset, shows the resulting averaged intensity profile. Corresponding distribution of the average events duration and amplitudes are shown in the center and right plots respectively. Overall, this comparative investigation supports our hypothesis that the elementary Ca^2+^ events triggered by endogenous Aβ oligomers are similar to those generated by the synthetic Aβ42 oligomers and IP_3_, and involve the opening of IP3Rs [23, 24].

To further strength our hypothesis, we examined the effect of caffeine, known to act as a reversible antagonist of IP_3_Rs, on the Ca^2+^ fluxes triggered by endogenous Aβs oligomers. Moreover, the ability of caffeine to freely permeate the oocyte’s membrane and the absence of ryanodine receptors, make caffeine an ideal agent to inhibit IP_3_Rs in Xenopus oocyte [8, 17, 25]. In these experiments, the oocytes were peeled (the vitelline membrane was removed) to increase oocytes adhesion to the coverslip ensuring the recording at the same exact location during application and after the washing of caffeine from the bathing solution. In Figure 6A a set of three traces are shown, reporting fluorescent intensity recorded at the same location right after injection of 10 nl of AD3-TEC sample at the time indicated by the arrow. The initial spike after ∼5 second followed by a much slower change in fluorescent signal seen in this experiment, was often observed during this investigation. Perfusion of caffeine in the bathing solution was performed by gravity driven fashion with a flux of about 0.4 ml/sec, able to fully exchange the volume of the chamber with 10 sec. To ensure full concentration of caffeine in the recording chamber, perfusion was left effective for about 2 to 3 minutes and then stopped. After 5 minutes incubation with caffeine, with the injecting needle left unmoved, a second injection of 10 nl of AD3-TEC sample was performed and the corresponding recording of the fluorescent signal is reported in Figure 6A center trace, showing the ability of caffeine to completely abolish Ca^2+^ fluorescent transient triggered when AD3-TEC was injected alone. Confirming the reversibility of caffeine’s effect on the IP_3_Rs, injection of 10 nl of AD3-TEc completely restored the Ca^2+^ response after extensive washing-of caffeine from the bathing solution (right top trace). Similar results were obtained from experiments injecting 10 nl AD3-B11 into oocytes with and without caffeine (Figure 5B). Interestingly, in the experiment shown in Figure 6C, scattered local transient puffs, were visualized throughout the membrane patch. Application of caffeine completely abolished the activity at each puff site. Puffs activity was restored after removal of caffeine as shown in Figure 6C right panel where 10 nl of AD1-B11 sample was injected at the same location.

**Figure 6).**
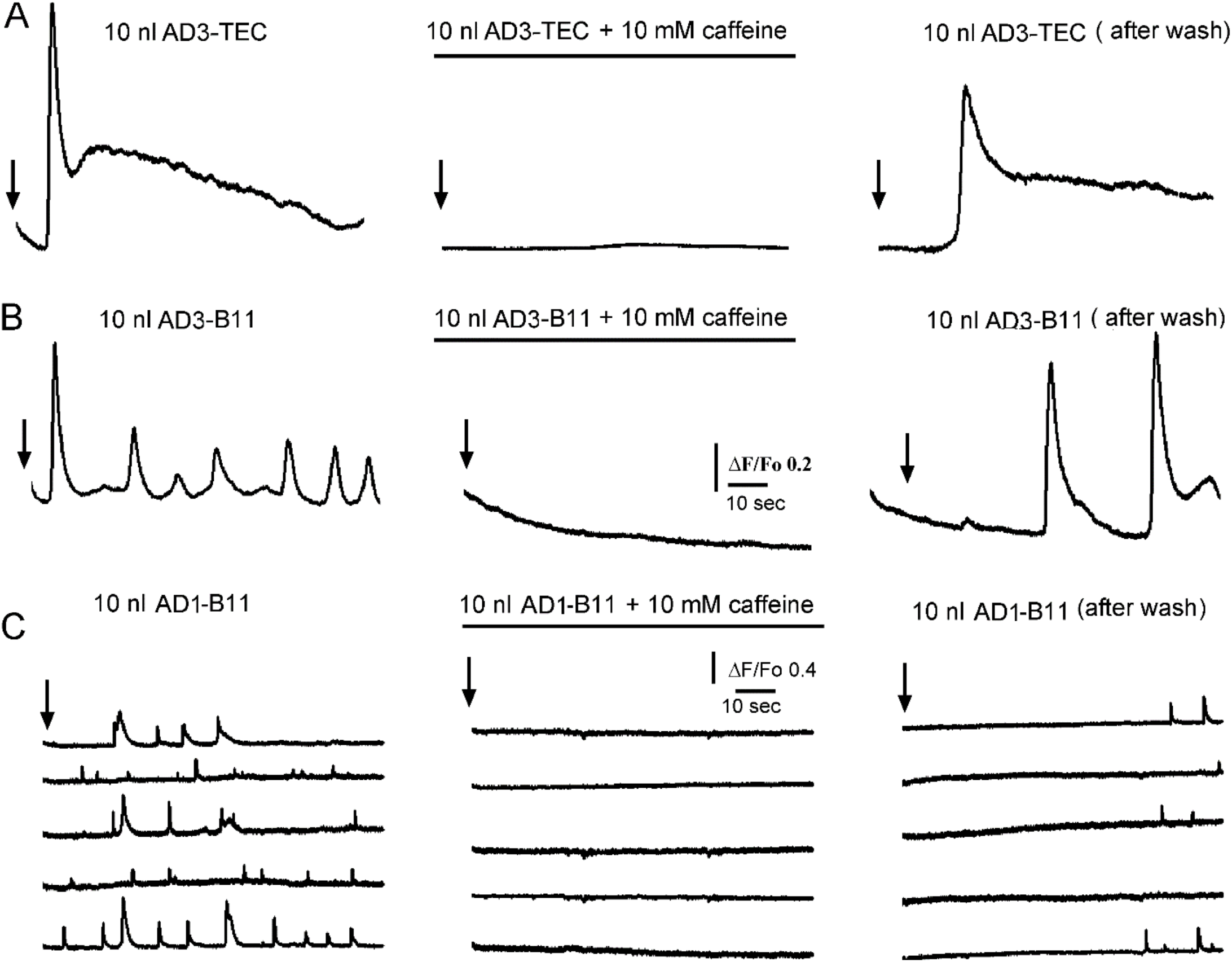
Caffeine reversibly inhibits local and global Ca^2+^ signaling triggered by intracellular injection of brain extracts. **A**) Left trace showing global Ca^2+^ fluorescent signal triggered by injection of 10 nl of AD3-TEC. Black arrows, indicate time of injection in each experiment. Central panel show equivalent experiment in the presence of caffeine, and right panel after caffeine washout in the same oocyte. **B and C**) are the experiments performed in different oocytes during intracellular injection of 10 nl of AD1-B11 and AD3-B11 brain extracts respectively. Signals in **A** and **B** are recorded from 40 × 40 μm regions of interest (global), whereas traces in **C** are from 3×3 pixels (1 μm^2^) regions centered on 5 different event sites within the imaged field.

### Computational quantification of the Ca^2+^ released and the corresponding amount of IP3 overproduction by cytosolic injection brain extracts

To quantify the amount of IP3 generated and consequently the amount of Ca^2+^ released from intracellular stores in response to intracellular injection of brain extracts from normal and AD-affected brains, we fitted the whole-cell model discussed in the Methods section to the observed fluorescence traces. Sample fits to the fluorescence traces are shown in Figure 7A-D, where red lines represent the fluorescence changes estimated by the model. Observed fluorescence changes (black) are shown for comparison. The blue curves represent the change in cytosolic Ca^2+^ concentration in response to brain extracts injection. The time-traces for Ca^2+^ and IP_3_ concentrations are integrated to estimate the total Ca^2+^ and IP_3_ concentrations generated by brain extracts from control brain and three different AD brains, and are shown in Figure 7E and F, respectively. Figure 7E and F clearly show a significantly larger IP_3_ generated and consequently higher Ca^2+^ released from intracellular stores due to extracts from AD patients as compared to extracts from control brains.

**Figure 7).**
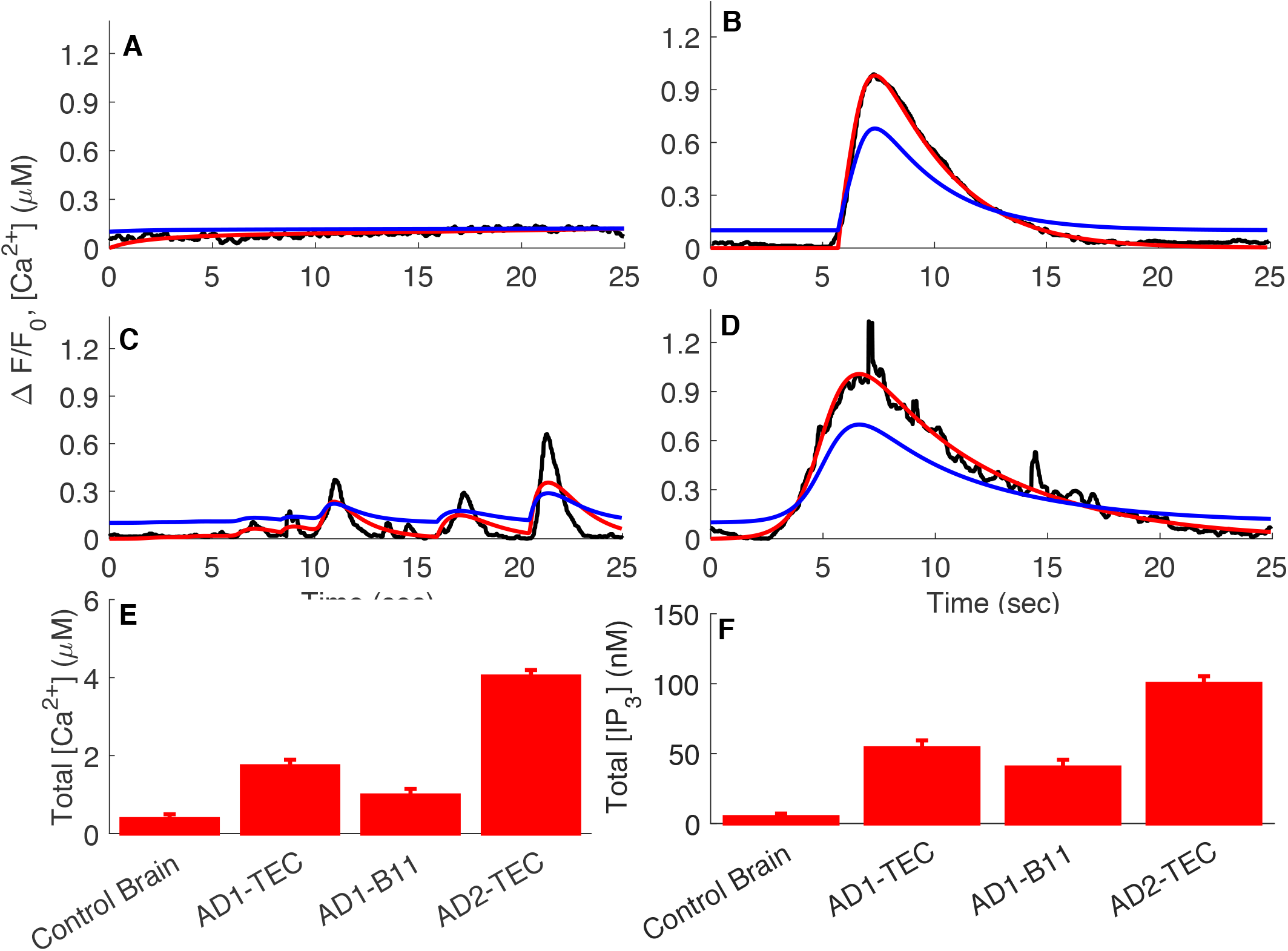
Computational analyses of the fluorescence signals allows quantification of amount of Ca^2+^ released and the corresponding IP_3_ generated by each brain extract. In Panels **A to D**, the black trace is ΔF/F0 (experiment), red is ΔF/F0 (model), and blue is the estimated Ca^2+^ from the model. **A, B, C, and D** are sample traces from Control brain, AD3-TEC, AD3-B11, and AD2-B11. The bar plots in **E and F** show the total amount of Ca^2+^ released and IP_3_ generated by brain extracts in 20 seconds, both estimated from the model fits. The error bars represent standard deviation from two experiments.

### Ca^2+^ dependent disruption of bioenergetics

Next, we used our model to estimate the bioenergetic cost of the aberrant Ca^2+^ rises due to brain extracts. Specifically, we feed the Ca^2+^ traces estimated above to a model for mitochondrial bioenergetics to estimate the changes in mitochondrial Ca^2+^ concentration, cell’s ATP, and ROS levels as detailed in the Methods section. Sample traces for these three variables in cells injected with control and AD2-TEC extracts are shown in Figure 8A, C, and E, respectively. A summary of changes in mitochondrial Ca^2+^ concentration, ATP and ROS obtained from fits to two experiments are shown Figure 8B, D, and F, respectively. Both the traces and bar plots reveal a significant drop in ATP and rise in ROS levels in cells injected with extracts from AD-affected brains.

**Figure 8).**
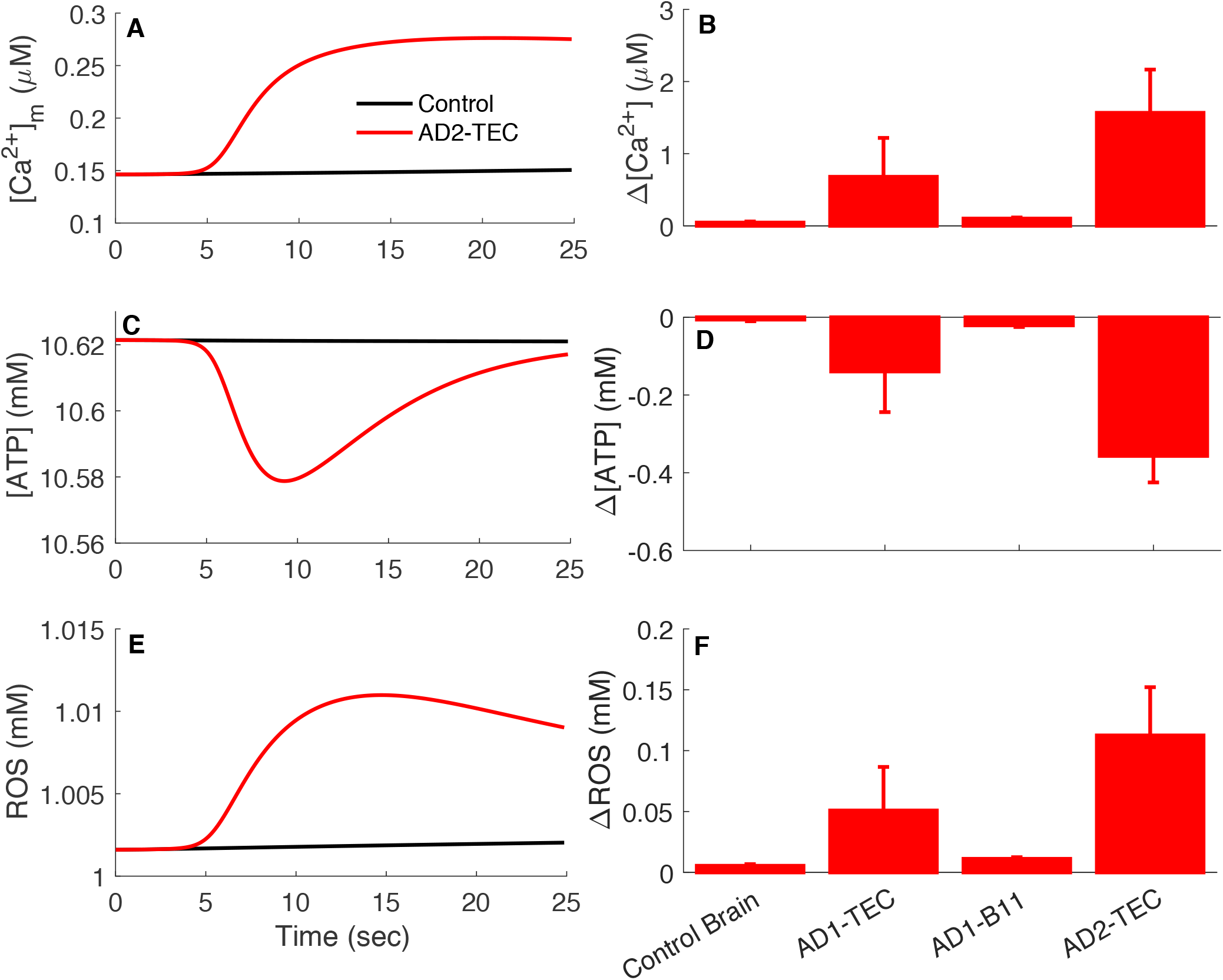
Rises in cytosolic Ca^2+^ due to AD brains extracts result in decreased ATP and higher ROS production as compared to extract from control brains. Panels **A, C, and E** display sample traces showing changes in mitochondrial Ca^2+^, cellular ATP, and ROS, respectively, in oocytes injected with extract from control (black) and AD-affected brains (red). The bar plots in **B, D, and F** show the total amount of Ca^2+^ buffered by mitochondria, decrease in ATP, and increase in ROS in 20 sec due to brain extracts. The error bars represent standard deviation from two experiments.

### Conclusions

A great deal of progress has been made in recent years to better understand the involvement of the amyloid beta peptide (Aβ) in the progression of Alzheimer’s disease [26-28]. A widely accepted hypothesis implicates the oligomeric forms of Aβs as the main toxic agent triggering an uncontrolled elevation of cytosolic Ca^2+^ to a poisonous level with consequent impairment on cell’s normal functioning and final death [1, 2, 7, 14, 29]. Using Ca^2+^-dependent fluorescent imaging, we revealed two distinct mechanisms by which Aβ42 affect intracellular Ca^2+^ homeostasis. One, by forming self-gating Ca^2+^-permeable pores allowing uncontrolled influx of extracellular Ca^2+^ into the cell’s cytosol, and a second mechanism involving the release of Ca^2+^ from intracellular stores through the opening of IP_3_R channels [8, 9]. These two mechanisms have been widely investigated by various groups using many different approaches [30-34]. Nevertheless, most of these investigations have been carry out using synthetic Aβs peptides, living some ambiguity whether the proposed mechanisms of action can be translated to the effect of endogenous Aβ oligomers found in neurons of AD-affected brains [35]. Here, we report the ability of brain extract samples, displaying high content of Aβ oligomers, to evoke cytosolic Ca^2+^ release upon intracellular injection, closely mirroring the effect of synthetic Aβ42 oligomers in analogous experiments [8]. The samples consist of homogenate extracts from postmortem brain tissues from frontal cortex from donors with clinical and pathological details shown in Table 1. Three samples from the frontal cortex (B11) of normal individual were used as control and compared with four samples (three B11 and one TEC) from three individuals exhibiting the typical hallmarks of AD-affected brains (Table 1).

The immunological characterization shown in Figure 1A reports dot blot analysis of fraction of samples probed with conformations specific OC and A11 antibodies and sequence specific 6E10 and 4G8 antibodies [36-40]. All the samples we tested, display low immunoreactivity to A11, 6E10 and 4G8, whereas high immunoreactivity was detected for OC antibody (Figure 1B). Quantitative analysis of the dot blot data shows significant level of OC immunoreactivity in the AD samples compared to normal subjects, in all the brain regions examined. Particularly, AD3 samples, for both B11 and TEC regions, display the higher content of OC-positive Aβ oligomers. Moreover, AD3 samples showed increased plaque stage as compared to AD1 and AD2 (Table 1). Intriguingly, all the AD samples display low immunoreactivity to A11, indicating that the endogenous Aβ oligomers contained in the brain samples have a specific aggregation state, equivalent to that acquired by the synthetic Aβ42 oligomers in our previous preparations [8, 9]. As shown in Figure 2, intracellular injections of 10 nl samples from AD3-TEC, AD3-B11 and AD1-B11 donors, all triggered cytosolic Ca^2+^ fluxes ranging from local events to global event involving large areas in the image field. Conversely, 10 nl N2-B11 sample from control brain, failed to trigger any significant response similarly to the effect of 10 nl injection of PBS (Figure 3D). The absence of Ca^2+^ in the bathing solution and the location of these Ca^2+^ events deep in the cytosol, point to a release of Ca^2+^ from intracellular stores. High variability in amplitudes and temporal evolution of the fluorescent signals, including local and global events evoked by all the AD affected samples (Figure 3A) closely resemble the fluorescent responses obtained after intracellular injection of synthetic Aβ42 oligomers and by injection of the agonist IP_3_ [8, 15].

Intracellular elementary events in the IP_3_ pathway signaling are the building blocks by which local and global Ca^2+^ signaling are orchestrated [16]. The strong similarity of the elementary events evoked by AD brain extracts with those evoked by the direct injection of the agonist IP_3_, supports the hypothesis of Ca^2+^ release from intracellular stores through activation of IP_3_Rs. This proposition is reinforced by experiments where we challenge the ability of brain extracts to trigger Ca^2+^ events by pre-incubating oocytes with 10 mM caffeine (Figure 5). Caffeine acts as a reversible, membrane-permeant competitive antagonist at the IP_3_R [8, 17, 25]. The specific involvement of the endogenous Aβ oligomers as the agent responsible for the Ca^2+^ mobilization by the brain extract is further strength by the ability of conformation-specific antibody OC to specifically decrease or reverse the ability all the brain extract to trigger Ca^2+^ fluxes (Figure 4). The fact that neither the sequences specific 6E10 nor 4G8 antibody display any detectable immunoreactivity in all of the AD extract suggest that most of the Aβs present in the samples are in their respective aggregated forms, and not in random coil monomer [18]. Moreover, the finding that among the two non-sequence specific/conformation specific antibody, only OC but not A11 showed high immunoreactivity in all the samples suggest that a specific conformation OC-positive Aβ oligomer is present in substantial amount in the cortex of AD affected brains [4, 5, 18].

Using pharmacological and computational investigation, we proposed that one of the mechanisms by which synthetic Aβ42 oligomers affect cells functioning involve the overproduction IP_3_ through stimulation of the G-protein complex triggering uncontrolled release of Ca^2+^ though the IP_3_Rs [8, 10, 41]. Intriguingly, these experiments were carried out using synthetic Aβ42 oligomers with high immunoreactivity to OC antibody, implying that similar toxicity may be carry out by endogenous intracellular Aβ oligomer on neurons of AD affected brains. Interesting, the ability of OC to react with Aβ aggregates that range from a few aggregated peptides to up to 40-50 peptides suggests that more than the sized of the aggregate is their specific aggregation. These results, supports the hypothesis that the intracellular rise of endogenous amyloid oligomer observed in neurons of AD-affected brains represent the toxic agents responsible for neurons malfunctioning and death, associated with the disruption of neuronal Ca^2+^ homeostasis.

To investigate the downstream effects of the aberrant cytosolic Ca^2+^ rises from the ER due to extracts from AD-affected brains, we first estimated the amount of IP_3_ generated and Ca^2+^ released by fitting a computational model of Ca^2+^ homeostasis to the observed florescence traces. Next, the time-traces of Ca^2+^ concentration were merged with a model for cell’s bioenergetics to assess the ATP consumed and ROS produced. Our simulations show a significant ATP loss and ROS rise due to extracts from AD-affected brains only in short duration. We believe that over periods, the pathological Ca^2+^ signals and consequently the low ATP and high ROS levels would make the cell progressively vulnerable, eventually leading to the cell dysfunction and demise. Overall, we believe that our study makes a strong case for disruptions in intracellular Ca^2+^homeostasis and consequently in cell’s bioenergetics as possible mechanism for the toxicity of endogenous Aβ oligomers.

## Supporting information

Supplementary Information Text: Computational Methods

## Acknowledgments

This work was supported by NIH grants R01AG053988 (to GU and AD), R01GM065830 (to IP), and RF1AG056507-01 (to CG).

