## Supplementary Information Text: Computational Methods for "Intracellular injection of brain extracts from Alzheimer’s disease patients trigger unregulated Ca^2+^ release from intracellular stores that hinders cellular bioenergetics"

### *Whole-cell $Ca^{2+}$ signaling model*

Extracts from AD-affected brains stimulate the production of $IP_3$ through PLC, which binds to $IP_3Rs$ to release $Ca^{2+}$ from the ER ($J_{ipr}$). $Ca^{2+}$ is also released from the ER through leak channels ($J_{leak}$), pumped back into the ER through Sarco/ER $Ca^{2+}$-ATPase (SERCA) ($J_s$), and buffered by $Ca^{2+}$ sensitive dye Fluo-4 ([dye]). Thus the rate equation for cytosolic $Ca^{2+}$ concentration ($[Ca^{2+}]_i$) is given as

$$\frac{d[Ca^{2+}]_i}{dt} = J_{ipr} + J_{leak} - J_s + k^r_{dye}\left(B_{dye} - [dye]\right) - k^f_{dye}[Ca^{2+}]_i[dye].$$

Where $B_{dye}$ is the total concentration of Fluo-4 and $k^f_{dye}$ and $k^r_{dye}$ are the binding and unbinding rates of $Ca^{2+}$ to Fluo-4 respectively. The functional form of the three fluxes are adopted from [1] and given as

$$J_{ipr} = k_{ipr}O([Ca^{2+}]_{ER} - [Ca^{2+}]_i),$$

$$J_{leak} = k_{leak}([Ca^{2+}]_{ER} - [Ca^{2+}]_i),$$

$$J_s = \frac{V_s[Ca^{2+}]_i^{ns}}{K_s^{ns} + [Ca^{2+}]_i^{ns}}.$$

$Ca^{2+}$ concentration in the ER ($[Ca^{2+}]_{ER}$) is given by the conservation of total $Ca^{2+}$ ($[Ca^{2+}]_t$) in the cell, i.e.

$$[Ca^{2+}]_{ER} = \gamma([Ca^{2+}]_t - [Ca^{2+}]_i).$$

Rate equation for Fluo-4 is given as

$$\frac{d[dye]}{dt} = k^r_{dye}\left(B_{dye} - [dye]\right) - k^f_{dye}[Ca^{2+}]_i[dye].$$

Parameter values used in the whole-cell $Ca^{2+}$ model are given in [2].

To model the gating of $IP_3R$, we use our previously developed kinetic scheme that closely replicates the behavior of type 1 $IP_3R$ in Xenopus Laevis oocytes (see Fig. 2 in [3]). The model has four states: a rest state (*R*) with no $Ca^{2+}$ bound, an active state (*A*) with 2 $Ca^{2+}$ bound, an open state (*O*) with 2 $Ca^{2+}$ bound, and an inhibited state (*I*) with 5 $Ca^{2+}$ bound. The rate equations for these four states are given as

$$\frac{dA}{dt} = K_{RA}R + K_{OA}O - (K_{AR} + K_{AO})A,$$

$$\frac{dO}{dt} = K_{IO}I + K_{AO}A - (K_{OI} + K_{OA})O,$$

$$\frac{dI}{dt} = K_{RI}R + K_{OI}O - (K_{IO} + K_{IR})I.$$

The fraction of channels in state *R* is given by the conservation of total probability, i.e.

$$R = 1 - A - O - I.$$

$K_{XY}$ represents the transition rate from state $X$ to $Y$. Various transition rates are given as

$$K_{RA} = \left[KR\left(\frac{1}{k_{01}[Ca^{2+}]_i} + \frac{1}{k_{12}[Ca^{2+}]_i^2}\right)\right]^{-1},$$

$$K_{AR} = \left[KA[Ca^{2+}]_i^2\left(\frac{1}{k_{01}[Ca^{2+}]_i} + \frac{1}{k_{12}[Ca^{2+}]_i^2}\right)\right]^{-1},$$

$$K_{AO} = \frac{k_{22}}{KA},$$

$$K_{OA} = \frac{k_{22}}{KO},$$

$$K_{OI} = \left[KO[Ca^{2+}]_i^2\left(\frac{1}{k_{23}[Ca^{2+}]_i^3} + \frac{1}{k_{45}[Ca^{2+}]_i^5}\right)\right]^{-1},$$

$$K_{IO} = \left[KI[Ca^{2+}]_i^5\left(\frac{1}{k_{23}[Ca^{2+}]_i^3} + \frac{1}{k_{45}[Ca^{2+}]_i^5}\right)\right]^{-1},$$

$$K_{RI} = \left[KR\left(\frac{1}{\tilde{k}_{01}[Ca^{2+}]_i} + \frac{1}{\tilde{k}_{45}[Ca^{2+}]_i^5}\right)\right]^{-1},$$

$$K_{IR} = \left[KI[Ca^{2+}]_i^5\left(\frac{1}{\tilde{k}_{01}[Ca^{2+}]_i} + \frac{1}{\tilde{k}_{45}[Ca^{2+}]_i^5}\right)\right]^{-1}.$$

*KR*, *KA*, *KO*, and *KI* are the occupancy parameters of state *R*, *A*, *O*, and *I* respectively with $KR = 1$ and

$$KA = \frac{a_1 a_3^{a_2}}{[IP_3]^{a_2} + a_3^{a_2}} + a_4,$$

$$KI = \frac{a_5 a_7^{a_6}}{[IP_3]^{a_6} + a_7^{a_6}} + a_8,$$

$$KO = \frac{a_9 [IP_3]^{a_{10}}}{[IP_3]^{a_{10}} + a_{11}^{a_{10}}}.$$

Various constant involved in the $IP_3R$ model are given in [2].

#### *Converting $Ca^{2+}$-bound dye to fluorescence units*

TIRF microscope measures changes in $[Ca^{2+}]_i$ in terms of fluorescence changes ($\Delta F/F_0$) as $Ca^{2+}$ binds and unbinds to indicator dye Fluo-4. Where $F_0$ and $\Delta F$ is background fluorescence and change in fluorescence in response to $Ca^{2+}$ binding to Fluo-4. Thus, we convert $Ca^{2+}$-bound dye from concentration units to $\Delta F/F_0$, which was then used in fitting the model to fluorescence signals from TIRF microscopy.

A single channel opening on average increases the fluorescence of a 1.2μm × 1.2μm area around the channel relative to the background signal by 0.11 ± 0.01 (i.e. $\Delta F/F_0$ = 0.11 ± 0.01) in TIRF microscopy experiments using Fluo-4 [4]. We used this information to convert $[Ca^{2+}]$-bound [dye] to $\Delta F/F_0$ in the model. We simulate a single channel, placed at the center of a 40μm × 40μm area (equal to the area scanned in the TIRF microscopy experiments in [4] as well as in our experiments) and allow it to open for 20ms (the mean open time of $IP_3R$). The $[Ca^{2+}]_i$ and [dye] equations described above are modified to include diffusion of these two species with the widely accepted diffusion coefficients of 223μm$^2$/s and 200μm$^2$/s for free $Ca^{2+}$ ($D_{Ca}$) and Fluo-4 ($D_{dye}$) respectively. Furthermore, to side-step the requirement of using extremely small spatial grid and stay consistent with the spatial resolution of 0.3μm of the TIRF microscopy experiments in [4], we partially adopt the procedure in [1], which simulates the $Ca^{2+}$ release and uptake at the microdomain around the channel separately from the rest of the simulation area. The movement of $Ca^{2+}$ from the microdomain to the nearest gird point (the central grid point in our case) and vice versa is given by diffusion. With these modifications, the equation for $[Ca^{2+}]_i$ becomes

$$\frac{d[Ca^{2+}]_i}{dt} = D_{Ca}\nabla^2[Ca^{2+}]_i + J_{diff} + J_{ipr} + J_{leak} - J_s + k^r_{dye}(B_{dye} - [dye]) - k^f_{dye}[Ca^{2+}]_i[dye].$$

Where the first and second terms represent the diffusion of $Ca^{2+}$ between neighboring grid points and the transfer of $Ca^{2+}$ from the microdomain around $IP_3R$ to the grid point at the center of simulating area respectively. The functional form of $J_{diff}$ is adopted from [1] where it is modeled with a function similar to Fick's first law, that is, $J_{diff} = k_{diff}([Ca^{2+}]_b - [Ca^{2+}]_i)$. For all other grid points, $J_{diff} = 0$. $[Ca^{2+}]_b$ is the $Ca^{2+}$ concentration in the microdomain and is given by the rate equation

$$\frac{d[Ca^{2+}]_b}{dt} = \gamma_1(J_{ipr} - J_{diff}).$$

$J_{ipr}$ for the single channel is governed by the current through the channel, i.e.

$$J_{ipr} = \frac{1}{2\times F\times\delta V}.$$

Where $I = 0.05pA$ is the observed current through $IP_3R$ [5], F is Faraday's constant, and δVis the volume of a hemisphere over the channel with radius of 10nm [6, 7].

With the inclusion of the microdomain in the model, the expression for $[Ca^{2+}]_{ER}$ changes accordingly:

$$[Ca^{2+}]_{ER} = \gamma([Ca^{2+}]_t - [Ca^{2+}]_i\text{-}[Ca^{2+}]_b/\gamma_1).$$

With the inclusion of diffusion, the rate equation for [*dye*] changes to

$$\frac{d[dye]}{dt} = D_{dye}\nabla^2[dye] + k^r_{dye}(B_{dye} - [dye]) - k^f_{dye}[Ca^{2+}]_i[dye].$$

The above equations were simulated using forward difference method and the peak change in $Ca^{2+}$-bound dye ($\Delta[dyeCa^{2+}]$) with respect to resting level ($[dyeCa^{2+}]_0$) was recorded, where $[dyeCa^{2+}] = B_{dye} -$ [dye]. The ratio $\Delta[dyeCa^{2+}]/[dyeCa^{2+}]_0$ averaged over 1.2μm × 1.2μm area around the channel together with the experimentally observed mean $\Delta F/F_0$ during a single $IP_3R$ opening was used to convert $Ca^{2+}$-bound dye to $\Delta F/F_0$.

***Estimating $IP_3$ production due to brain extracts***

Injection of brain extracts into the oocytes stimulates $IP_3$ production through a G-protein coupled mechanism. $IP_3$ activates $IP_3R$ on the ER membrane, releasing $Ca^{2+}$ into the cytoplasm. The global $[Ca^{2+}]_i$ is then represented by $\Delta F/F_0$, imaged through TIRF microscopy. As shown in the main text, after injecting brain extracts, oocytes show a range of fluorescence responses. Guided by the time-trace representing the average fluorescence response in a cell, we write an arbitrary function for $[IP_3]$ ($f[IP_3]$) that loosely resembles the fluorescence traces. For example, for the trace in Fig. 7B (main text), we choose the function $f[IP_3]$ to be

$$f[IP_3] = p_1\left[\frac{1}{1+exp\left(\frac{p_2-t}{p_3}\right)}\right]exp\left(-\frac{t}{p_4}\right),$$

Where $t$ is time in seconds starting from the injection of brain extract and $p_1 - p_4$ are arbitrary parameters.

Next, the whole-cell model together with $f[IP_3]$ and $Ca^{2+}$-bound [dye] to $\Delta F/F_O$ conversion) is fitted to the fluorescence trace, optimizing the parameters in $f[IP_3]$ so that the whole-cell $Ca^{2+}$ signaling model gives the best fit to the fluorescence traces. The optimizing in performed in Matlab by computing the least squares error ($\chi^2$) as follows.

The Matlab code first solves the rate equations for the whole-cell and uses the conversion factor for $Ca^{2+}$-bound dye to $\Delta F/F_O$ to get the whole-cell fluorescence signal given by the model ($\Delta F/F_{O,M}$). Using the whole-cell fluorescence signal from TIRF microscopy experiments ($\Delta F/F_{O,E}$) as observable, we can write $\chi^2$ as

$$\chi^2 = \frac{1}{N}\sum_{k=1}^{N}\left[\left(\frac{\Delta F}{F_{o,M}}\right)_k - \left(\frac{\Delta F}{F_{o,E}}\right)_k\right]^2,$$

where $N$ is the number of data points in the experimental traces. Various parameters in $f[IP_3]$) are determined by minimizing $\chi^2$ for whole-cell TIRF signals. The same procedure is repeated for all oocytes with extracts from different brains.

#### *Mitochondrial function model*

The rate equations modeling mitochondrial function are adopted from our previous work [2, 8, 9] and are described in [2] in detail. These equations are originally based on the model in [10]. The model includes processes such as the tricarboxylic acid cycle, electron transport chain, $Ca^{2+}$ signaling pathways, and reactive oxygen species (ROS) production. The time traces of cytosolic $Ca^{2+}$ estimated above are coupled with the mitochondrial model through the rate equation for mitochondrial $Ca^{2+}$ concentration ($[Ca^{2+}]_m$) to investigate changes in cell's ATP and ROS levels due to brain extracts injection. Increase in $[Ca^{2+}]_m$ stimulates the production of ATP through the tricarboxylic acid cycle, decreases ATP production by depolarizing mitochondrial membrane, and increases ROS production through electron transport chain. Similarly, increase in cytosolic $Ca^{2+}$ causes a decrease in cell's ATP as base cytosolic $Ca^{2+}$ level is restored by ATP-consuming processes.

#### *Numerical and Experimental Methods*

Simulations in section "Converting $Ca^{2+}$-bound dye to fluorescence units" were performed using forward difference method with a time-step of 50μs and a spatial grid size of 0.3μm, equal to the pixel size in the experiments in [4] that estimated the mean $\Delta F/F_O$ value during a single channel opening. Reducing the grid size and time-step did not change our estimates significantly. The initial value of $[Ca^{2+}]_i$ was set to 50nM, approximately equal to the resting cytosolic $Ca^{2+}$ concentration. Based on our simulations, we observed that in resting state approximately 1μM of fluorescence dye was bound to $Ca^{2+}$. Thus, we set the initial value of [dye] = 39μM. The boundaries were fixed accordingly at steady state values of $[Ca^{2+}]_i$ = 50nM and [dye] = 39μM.

Since there was no stimulus at the start of the experiments, $IP_3Rs$ were initially considered to be in the resting state, that is, the initial values of R, A, O, and I were set at 1, 0, 0, and 0 respectively. In all cases, simulations were allowed to reach steady state before applying any stimulus, so selecting slightly different initial conditions would not change the final results.

To fit the whole-cell model to the TIRF signals, the rate equations were solved in Matlab using the in-built function “ode15s”. $\chi^2$ function was minimized using the in-built Matlab function “fminsearch”. Using other Matlab optimization functions such as “lsqcurvefit”, ”fmincon”, or “nlinfit” did not change the quality of fits.

Numerical integration of the full model equations (cytosolic $Ca^{2+}$ dynamics, IP3Rs, mitochondrial $Ca^{2+}$ dynamics and bioenergetics) was performed with Intel Fortran compiler (Intel Corporation, Santa Clara, CA). ODEs were solved using RK4 method. Code producing key results in the paper is available upon request from the authors.
